# An engineered ACE2 decoy receptor can be administered by inhalation and potently targets the BA.1 and BA.2 omicron variants of SARS-CoV-2

**DOI:** 10.1101/2022.03.28.486075

**Authors:** Lianghui Zhang, Krishna K. Narayanan, Laura Cooper, Kui K. Chan, Christine A. Devlin, Aaron Aguhob, Kristie Shirley, Lijun Rong, Jalees Rehman, Asrar B. Malik, Erik Procko

## Abstract

Monoclonal antibodies targeting the SARS-CoV-2 spike (S) glycoprotein neutralize infection and are efficacious for the treatment of mild-to-moderate COVID-19. However, SARS-CoV-2 variants have emerged that partially or fully escape monoclonal antibodies in clinical use. Notably, the BA.2 sublineage of B.1.1.529/omicron escapes nearly all monoclonal antibodies currently authorized for therapeutic treatment of COVID-19. Decoy receptors, which are based on soluble forms of the host entry receptor ACE2, are an alternative strategy that broadly bind and block S from SARS-CoV-2 variants and related betacoronaviruses. The high-affinity and catalytically active decoy sACE2_2_.v2.4-IgG1 was previously shown to be effective in vivo against SARS-CoV-2 variants when administered intravenously. Here, the inhalation of sACE2_2_.v2.4-IgG1 is found to increase survival and ameliorate lung injury in K18-hACE2 transgenic mice inoculated with a lethal dose of the virulent P.1/gamma virus. Loss of catalytic activity reduced the decoy’s therapeutic efficacy supporting dual mechanisms of action: direct blocking of viral S and turnover of ACE2 substrates associated with lung injury and inflammation. Binding of sACE2_2_.v2.4-IgG1 remained tight to S of BA.1 omicron, despite BA.1 omicron having extensive mutations, and binding exceeded that of four monoclonal antibodies approved for clinical use. BA.1 pseudovirus and authentic virus were neutralized at picomolar concentrations. Finally, tight binding was maintained against S from the BA.2 omicron sublineage, which differs from S of BA.1 by 26 mutations. Overall, the therapeutic potential of sACE2_2_.v2.4-IgG1 is further confirmed by inhalation route and broad neutralization potency persists against increasingly divergent SARS-CoV-2 variants.

## INTRODUCTION

Monoclonal antibodies targeting the Spike (S) of severe acute respiratory syndrome coronavirus 2 (SARS-CoV-2) are clinically effective at reducing or preventing COVID-19 symptoms (Gupta *et al*, 2021; Weinreich *et al*, 2021; O’Brien *et al*, 2022; Gottlieb *et al*, 2021). As of March, 2022, six antibodies have received emergency use authorization from the U.S. Food and Drug Administration for treating mild-to-moderate COVID-19 (REGN10933/casirivimab, REGN10987/imdevimab (Hansen *et al*, 2020), LY-CoV555/bamlanivimab (Jones *et al*, 2021), LY-CoV016/etesevimab (Shi *et al*, 2020), VIR-7831/sotrovimab (Pinto *et al*, 2020), and most recently LY-CoV1404/bebtelovimab (Westendorf *et al*, 2022)) and another two antibodies have authorization for prophylactic administration as a slow-release cocktail in immunocompromised patients (AZD8895/tixagevimab and AZD1061/cilgavimab (Zost *et al*, 2020)). All authorized antibodies target the receptor-binding domain (RBD) of the S protein to neutralize infection. While the RBD has the conserved function of binding the human receptor for SARS-CoV-2 cell entry, the RBD sequence is itself poorly conserved across SARS-related betacoronaviruses (Chan *et al*, 2021). Mutational scans have demonstrated that many mutations are tolerated (Chan *et al*, 2021; Starr *et al*, 2020) and the RBD is a region of substantial diversity among SARS-CoV-2 variants in circulation (Hirabara *et al*, 2022). Mutations within the RBD allow immune escape and increase transmissibility via enhanced receptor affinity. Rapid viral evolution has been observed after treatment with monoclonal antibody drugs, including the appearance of escape mutations to LY-CoV555 and VIR-7831 in immunocompromised (Jensen *et al*, 2021) and immunocompetent patients (Rockett *et al*, 2022). To minimize the likelihood of full escape, non-competing monoclonal antibodies are combined as cocktails with some success (Baum *et al*, 2020). For example, the virulent P.1/gamma variant of concern (VOC) carrying 3 mutations in the RBD compared to original virus isolates is resistant to REGN10933 neutralization but is sensitive to REGN10987; the cocktail of the two antibodies remained effective (Copin *et al*, 2021).

The emergence and rapid spread of the B.1.1.529/omicron VOC has upended the development of monoclonal antibodies for COVID-19. The BA.1 omicron sublineage was first detected in southern Africa in November 2021 and rapidly spread within weeks to displace B.1.617.2/delta as the most prevalent VOC (Wang & Cheng, 2022; Viana *et al*, 2022). A second omicron sublineage, BA.2, has been steadily rising and is the dominant variant in some geographical regions (Lyngse *et al*, 2022). Omicron far exceeds other VOCs in its number of mutations; S proteins of BA.1 and BA.2 omicron have approximately 37 and 31 mutations compared to the original virus, of which only 21 mutations are shared by both sublineages (Majumdar & Sarkar, 2022). The RBDs alone, which are targeted by many antibodies, have 15 and 16 mutations, respectively, of which 12 are shared and many are localized to the receptor binding interface. Consequently, there are extensive changes to antigenic epitopes on the surface of S. Neutralizing antibody titers are diminished in the serum of recovered and vaccinated individuals (Planas *et al*, 2022; Cele *et al*, 2021; Rössler *et al*, 2022; Ikemura *et al*, 2022), and BA.1 omicron is reported to escape the REGN10933+REGN10987, LY-CoV555+LY-CoV016, and AZD1061+AZD8895 cocktails (Cao *et al*, 2022; VanBlargan *et al*, 2022; Planas *et al*, 2022; Ikemura *et al*, 2022). Very recently, it has been reported that VIR-7831 has markedly reduced efficacy against pseudovirus expressing S of BA.2 omicron, raising the possibility that only one authorized antibody (LY-CoV1404) may remain clinically effective (Zhou *et al*, 2022; Iketani *et al*, 2022). Indeed, in one study, just 2 antibodies from a panel of 19 in preclinical and clinical development remained potent against BA.2 omicron (Iketani *et al*, 2022). These findings challenge whether monoclonal antibodies are suitable over the long term for the treatment of endemic COVID-19.

An alternative approach is to use soluble decoy receptors that bind and block the RBD (Monteil *et al*, 2020; Jing & Procko, 2021; Chan *et al*, 2020; Hofmann *et al*, 2004; Lei *et al*, 2020). S binds to angiotensin-converting enzyme 2 (ACE2), which is highly expressed on type II alveolar lung epithelium (Hamming *et al*, 2004; Zhao *et al*, 2020), triggering conformational changes that facilitate fusion of the viral envelope and host cell membrane (Huang *et al*, 2020). ACE2 is a protease of the renin-angiotensin-aldosterone system (RAAS), which catalyzes the cleavage and inactivation of the vasoconstrictor angiotensin II (Ang-II) (Vickers *et al*, 2002; Tipnis *et al*, 2000), as well as the cleavage of other pro-inflammatory peptides in the kinin system that promote vascular leakage (Vickers *et al*, 2002), and formation of Ang 1-7 to mediate anti-inflammation, anti-fibrosis, and vasodilation through Mas signaling (Kuba *et al*, 2021; Santos *et al*, 2003). The extracellular domains of ACE2 can be expressed as a soluble protein (sACE2) that blocks the RBD (Hofmann *et al*, 2004). Recombinant sACE2 has been evaluated in hospitalized COVID-19 patients, where it was found to decrease time on mechanical ventilation but had no positive impact on survival (http://ClinicalTrials.gov Identifier NCT04335136). To improve efficacy, next generation sACE2 decoys have been engineered for exceptionally tight affinity to S (K_D_ < 1 nM), on par with monoclonal antibodies (Chan *et al*, 2020; Glasgow *et al*, 2020; Sims *et al*, 2021; Cohen-Dvashi *et al*, 2020; Higuchi *et al*, 2021).

ACE2-based decoys have two proposed advantages that distinguish them from antibodies. First, infection reduces ACE2 activity in the lungs due to cell death and shedding of cellular ACE2 through the action of proteases (Kuba *et al*, 2005; Haga *et al*, 2008; Heurich *et al*, 2014). This causes massive dysregulation of the RAAS, with large increases in serum Ang-II and serum sACE2 (Fagyas *et al*, 2022; Kragstrup *et al*, 2021; Filbin *et al*, 2021; Lundström *et al*, 2021; Reindl-Schwaighofer *et al*, 2021; Wu *et al*, 2020; Liu *et al*, 2020), although much of the serum sACE2 may have low catalytic activity as well as low S affinity and avidity due to proteolysis within the ACE2 collectrin-like dimerization domain. These serum markers are highly correlated with disease severity and elevated Ang-II may contribute to vasoconstriction, thrombophilia, microthrombosis, and respiratory failure. Administering catalytically active sACE2 can dampen Ang-II and kinin signaling to reduce lung injury (Treml *et al*, 2010; Zou *et al*, 2014; Imai *et al*, 2005; Bastolla *et al*, 2022). However, many groups have argued that sACE2 therapeutics must be catalytically inactivated to prevent off-target toxicity (Tanaka *et al*, 2021; Lei *et al*, 2020; Iwanaga *et al*, 2020; Cohen-Dvashi *et al*, 2020; Glasgow *et al*, 2020; Sims *et al*, 2021; Chen *et al*, 2021; Higuchi *et al*, 2021).

The second proposed advantage is that receptor decoys will have unparalleled breadth against variants of S (Chan *et al*, 2021, 2020). If a mutant S protein has diminished affinity for the decoy, it will likely have diminished affinity for the native receptor and the virus will be attenuated. Engineered sACE2 decoys tightly bind diverse SARS-CoV-2 variants, as well as related betacoronaviruses from bats (Chan *et al*, 2021; Zhang *et al*, 2022; Higuchi *et al*, 2021; Yao *et al*, 2021). Potent neutralization persists against BA.1 omicron for at least some engineered decoys (Ikemura *et al*, 2022), but activity against BA.2 omicron has yet to be tested.

Here, we evaluate these two perceived advantages of an engineered sACE2 decoy, demonstrating that ACE2 catalytic activity contributes to therapeutic efficacy and tight binding persists against the divergent S proteins of BA.1 and BA.2 omicron.

## RESULTS

### sACE2_2_.v2.4-IgG1 inhalation alleviates lung injury and increases survival of gamma infected mice

The engineered decoy sACE2_2_.v2.4-IgG1 has three substitutions compared to wild type ACE2 that enhance affinity for S by over an order of magnitude, measured against S proteins from SARS-CoV-2 variants predating omicron (Chan *et al*, 2021, 2020; Zhang *et al*, 2022). Fusion to the Fc region of IgG1 increases serum stability and virus clearance (Chen *et al*, 2021). We recently found that intravenous (IV) administration of sACE2_2_.v2.4-IgG1 mitigates lung vascular endothelial injury and increases survival in K18-hACE2 transgenic mice infected with SARS-CoV-2 variants (Zhang *et al*, 2022). To further characterize the translational potential of sACE2_2_.v2.4-IgG1, the protein was nebulized and administered by inhalation to K18-hACE2 transgenic mice. Mice were inoculated with a lethal dose of virulent SARS-CoV-2 isolate/Japan/TY7-503/2021 (P.1/gamma variant) at 1×10^4^ plaque forming units (PFU) to induce severe lung injury. sACE2_2_.v2.4-IgG1 (7.5 ml at 8.3 mg/ml in PBS) was aerosolized by a nebulizer and delivered to the mice starting from 12 h post-inoculation. 3 doses were given 36 hours apart (**Figure 1A**) and aerosol delivery of PBS was applied as a control. The doses and inhalation interval of sACE2_2_.v2.4-IgG1 were based on previously published pharmacokinetic studies using inhalation and direct intratracheal delivery (Zhang *et al*, 2022). We estimate the inhaled dose of sACE2_2_.v2.4-IgG1 to the lungs to be ~0.5 mg/kg/dose.

**Figure 1.**
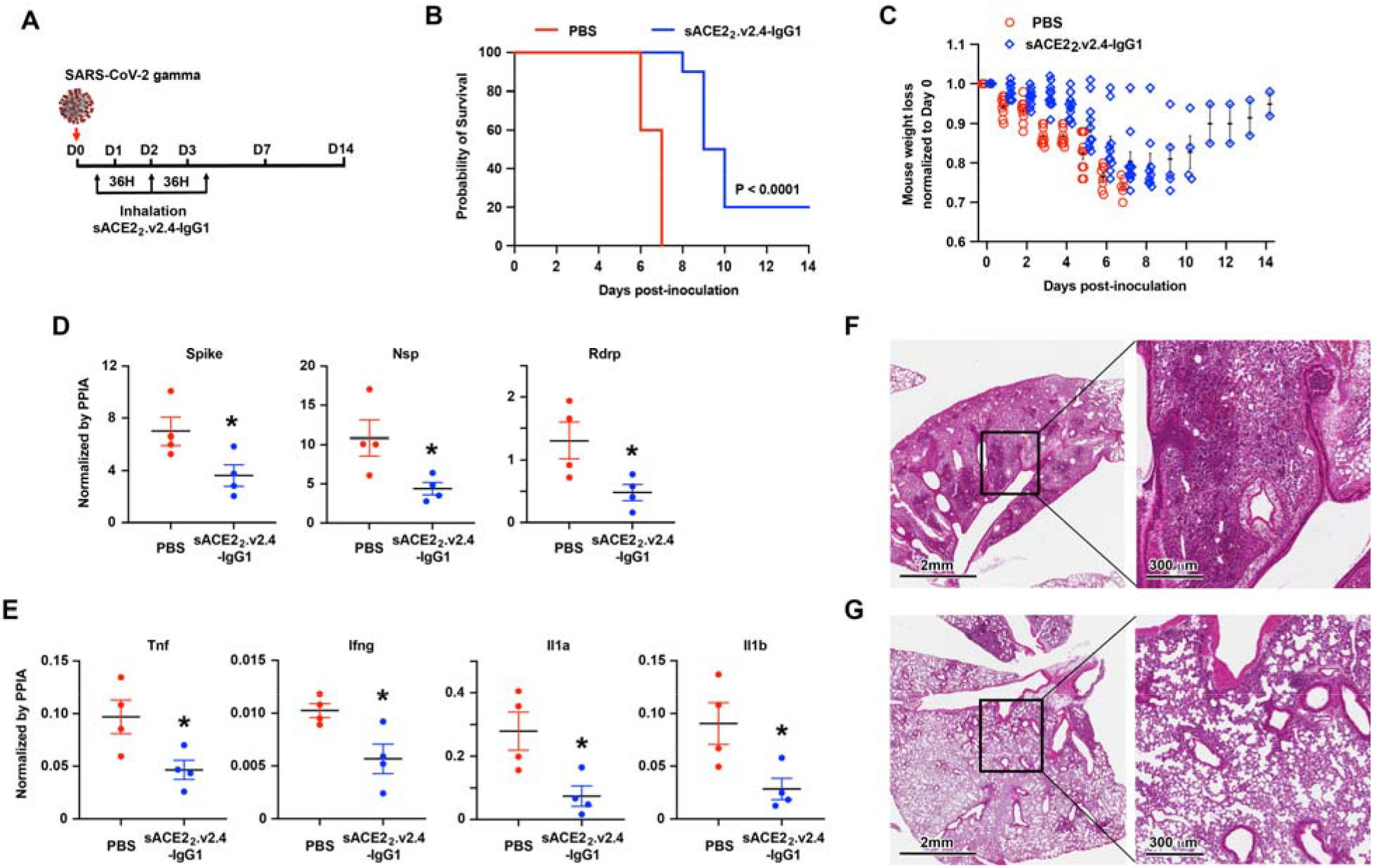
Aerosol delivery of sACE2_2_.v2.4-IgG1 alleviates lung injury and improves survival of SARS-CoV-2 gamma variant infected K18-hACE2 transgenic mice. **(A)** K18-hACE2 transgenic mice were inoculated with SARS-CoV-2 isolate /Japan/TY7-503/2021 (gamma variant) at 1×10^4^ PFU. sACE2_2_.v2.4-IgG1 (7.5 ml at 8.3 mg/ml in PBS) was delivered to the mice by a nebulizer in 25 minutes at 12 h, 48 h, and 84 h post-inoculation. PBS was aerosol delivered as control. **(B-C)** Survival curves **(B)** and weight loss **(C)**. N = 10 mice for each group. The P-value of survival curve by Gehan-Breslow-Wilcoxon test is shown. **(D)** Viral load in the lung was measured by RT-qPCR at Day 7. The mRNA expression levels of SARS-CoV-2 Spike, Nsp, and Rdrp are normalized to the house-keeping gene peptidylprolyl isomerase A (Ppia). **(E)** Cytokine expression levels of Tnf, Ifng, Il1a, and Il1b were measured by RT-qPCR normalized by Ppia. Data are presented as mean ± SEM. *, p < 0.05 by unpaired Student’s t-test with two sided. **(F-G)** Representative H&E staining of lung sections at Day 7 post-inoculation for control PBS group **(F)** and inhalation of sACE2_2_.v2.4-IgG1 group **(G)**. Images at left are low magnifications. Boxed regions (black) are shown at higher magnification on the right. Lungs from 4 independent mice were sectioned, stained, and imaged.

We chose the gamma variant to assess efficacy of the inhaled sACE2_2_.v2.4-IgG1 decoy because this variant has been shown to induce severe forms of COVID-19 resulting in high mortality (Lin *et al*, 2021). In our experimental model, all mice inoculated with the gamma variant and receiving PBS inhalation as a control died at 6-7 days (**Figure 1B**) with a 30% weight loss (**Figure 1C**). In the treatment group with inhalation of sACE2_2_.v2.4-IgG1, 20% of mice survived SARS-CoV-2 gamma infection with 5%-15% weight loss and 80% of mice died in 8-10 days, indicating prolonged survival (**Figures 1B and 1C**). Replication of SARS-CoV-2 gamma variant in the lung at Day 7 post-inoculation was measured by reverse transcription and real-time quantitative PCR (RT-qPCR) for the expression levels of SARS-CoV-2 Spike, non-structural protein (Nsp), and RNA-dependent RNA polymerase (Rdrp) (Figure 1D). We found that sACE2_2_.v2.4-IgG1 inhalation significantly inhibited viral replication in the lungs. Furthermore, we measured the expression levels of cytokines Tumor necrosis factor (Tnf), Interferon gamma (Ifng), Interleukin 1 alpha (Il1a), and Interleukin 1 beta (Il1b) in the lungs by RT-qPCR (**Figure 1E**). Aerosol delivery of sACE2_2_.v2.4-IgG1 reduced cytokine release in the lungs. Hematoxylin-eosin (H&E) staining of lung sections demonstrated severe immune cell infiltration induced by gamma variant infection at Day 7 in the control group (**Figure 1F**). sACE2_2_.v2.4-IgG1 inhalation significantly reduced immune cell infiltration at Day 7 (**Figure 1G**). Combining all, we conclude that sACE2_2_.v2.4-IgG1 inhalation efficiently alleviated severe lung injury and increased survival following a lethal dose of SARS-CoV-2 gamma variant infection by inhibiting viral replication and reducing cytokine release in the lung.

### Catalytic activity of sACE2_2_.v2.4-IgG1 contributes to therapeutic efficacy

To test whether proteolytic activity of a sACE2 decoy contributes to the mechanism of reducing disease following SARS-CoV-2 infection, two mutations (H374N and H378N, abbreviated “NN”) were introduced to the sACE2_2_.v2.4-IgG1 active site. These mutations disrupt the coordination site of an essential Zn^2+^ ion and were previously shown to have no impact on SARS-CoV-1 infection (Moore *et al*, 2004). We confirmed that catalytically dead sACE2_2_.v2.4(NN)-IgG1 failed to cleave a substrate peptide (**Figure 2A**), whereas the protein’s affinity for RBD was unchanged (**Figure 2B and 2C**).

**Figure 2.**
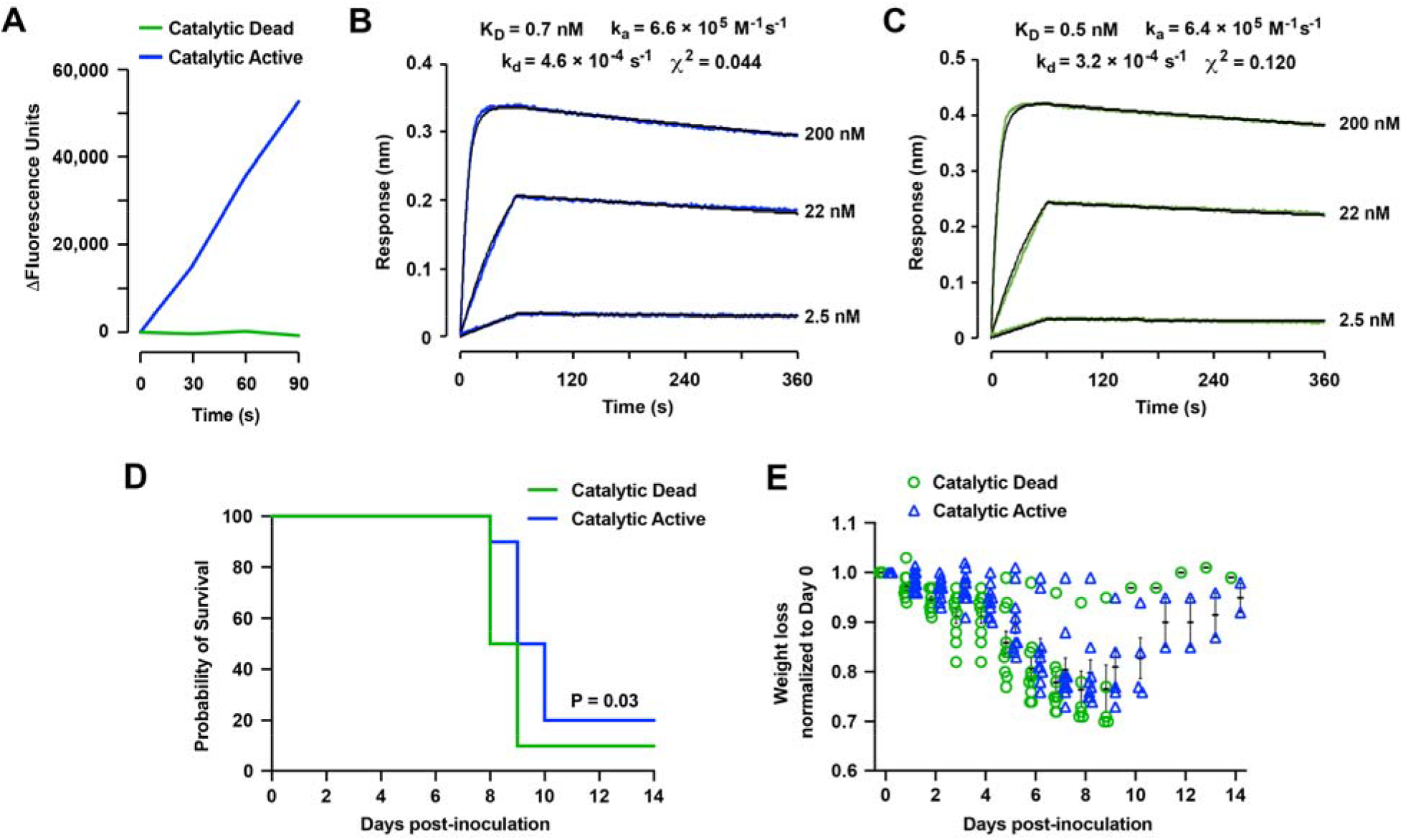
Catalytic activity of sACE2_2_.v2.4-IgG1 contributes to the therapeutic efficacy to mitigate mouse lung injury and improve survival following SARS-CoV-2 gamma infection. **(A)** A 7-Methoxycoumarin-4-acetyl (MCA) conjugated peptide is quenched by a 2,4-dinitrophenyl group. ACE2 catalyzed cleavage of the peptide is measured by increased MCA fluorescence. Mutations H374N and H378N generate a catalytically dead sACE2_2_.v2.4-IgG1 protein. **(B-C)** Catalytically active sACE2_2_.v2.4-IgG1 **(B)** and catalytically dead sACE2_2_.v2.4(NN)-IgG1 **(C)** were immobilized on BLI biosensors that were transferred to solutions of RBD as the soluble analyte (0-60 s) and returned to buffer to measure dissociation (60-240 s). RBD concentrations are indicated on the right of the sensorgrams. **(D-E)** Catalytically active sACE2_2_.v2.4-IgG1 and catalytically dead sACE2_2_.v2.4(NN)-IgG1 were aerosolized (7.5 ml protein at 8.3 mg/ml in 25 minutes) and delivered by inhalation to K18-hACE2 transgenic mice at 12 h, 48 h, and 84 h post-inoculation with SARS-CoV-2 gamma variant. 10 mice in each group were observed for survival **(D)** and weight loss **(E)**. The P-value of survival curve by Gehan-Breslow-Wilcoxon test is shown. Catalytically active and inactive proteins were tested in the same experiment versus PBS control shown in Figure 1.

We tested the therapeutic efficacy of catalytically dead sACE2_2_.v2.4(NN)-IgG1 directly against catalytically active sACE2_2_.v2.4-IgG1 using a lethal dose (1×10^4^ PFU) of SARS-CoV-2 gamma variant to infect K18-hACE2 transgenic mice. Consistent with its high affinity for blocking S and neutralizing infection, catalytically dead sACE2_2_.v2.4(NN)-IgG1 prolonged survival with 10% survival rate (**Figure 2D**). However, catalytically active sACE2_2_.v2.4-IgG1 extended survival further by ~1 day longer than catalytically dead sACE2_2_.v2.4(NN)-IgG1 with 20% survival rate (**Figure 2D**), supporting the hypothesis that ACE2 catalytic activity contributes to therapeutic efficacy. The mice that inhaled catalytically dead sACE2_2_.v2.4(NN)-IgG1 lost more body weight than mice that inhaled catalytically active sACE2_2_.v2.4-IgG1 (**Figure 2E**). This result agrees with seminal research demonstrating that ACE2 protects against lung injury (Treml *et al*, 2010; Imai *et al*, 2005; Zou *et al*, 2014) and is also supported by the observation that a bacterial ACE2 homologue protects SARS-CoV-2 infected animals, despite having no affinity for S (Yamaguchi *et al*, 2021). Overall, we conclude that sACE2_2_.v2.4-IgG1 has dual mechanisms of action: (i) blockade of receptor binding sites on SARS-CoV-2 spikes and (ii) turnover of vasoconstrictive and pro-inflammatory peptides that otherwise contribute to lung injury.

### sACE2_2_.v2.4-IgG1 tightly binds and neutralizes BA.1 omicron virus

Mature ACE2 is composed of a protease domain (amino acids [a.a.] 18-615) that contains the S interaction site, a collectrin-like dimerization domain (a.a. 616-732), a transmembrane domain (a.a. 741-762), and cytoplasmic tail (a.a. 763-805) (Yan *et al*, 2020). Soluble ACE2 from residues 18-615 is a monomeric protein and its binding to S-expressing cells is dependent on monovalent affinity. Soluble ACE2 from residues 18-732 is a stable dimer (which we denote as sACE2_2_) that binds avidly to S-expressing cells. Avid binding can mask differences in monovalent affinity (Zhang *et al*, 2022; Chan *et al*, 2020, 2021).

We incubated cells expressing BA.1 omicron S with three monomeric sACE2(18-615) proteins: wild type, v2.4 (ACE2 mutations T27Y, L79T, and N330Y), and a second engineered decoy called CDY14 (ACE2 mutations K31M, E35K, S47A, L79F, L91P, and N330Y; this is the highest affinity decoy reported in published literature (Sims *et al*, 2021)). Cells were washed and bound proteins were detected by flow cytometry. BA.1 omicron does not escape the engineered decoy receptors, and both sACE2.v2.4 and sACE2.CDY14 bind tighter than wild type sACE2 (**Figure 3A**). We note that sACE2.CDY14, with twice as many mutations, did not bind any tighter than sACE2.v2.4.

**Figure 3.**
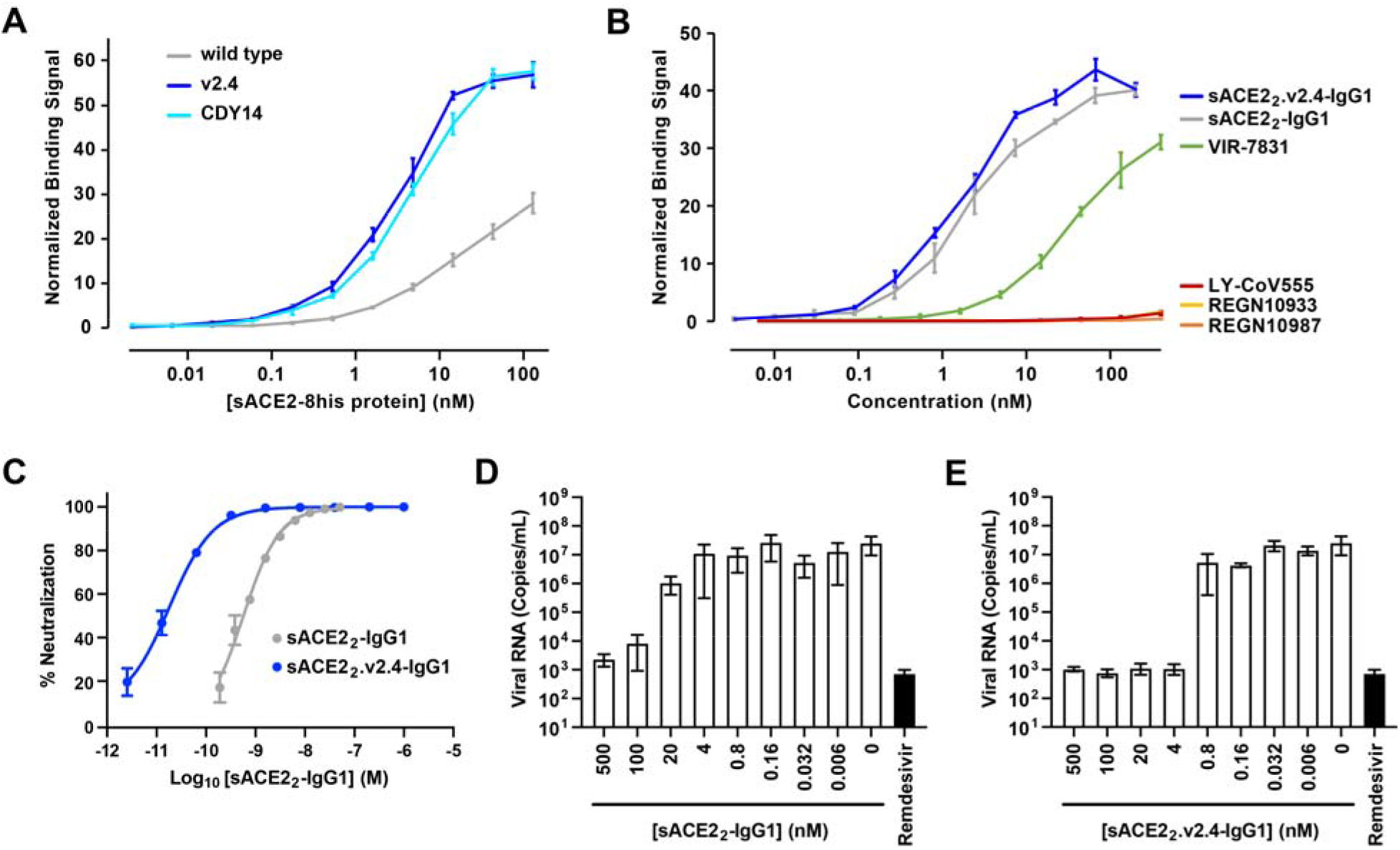
The engineered decoy tightly binds S of BA.1 omicron and neutralizes infection. **(A)** Binding of monomeric sACE2-8his proteins to Expi293F cells expressing S of BA.1 omicron was measured by flow cytometry. Data are mean ± SEM, N = 4 independent experiments. **(B)** Binding of dimeric sACE2_2_-IgG1 proteins was compared to the binding of antibodies authorized for therapeutic use in COVID-19 patients. Binding to Expi293F cells expressing BA.1 omicron S was measured by flow cytometry. Data are mean ± SEM, N = 4 independent experiments. **(C)** Neutralization of BA.1 omicron pseudovirus. sACE2_2_-IgG1 (grey) or sACE2_2_.v2.4-IgG1 (blue) were incubated with pseudovirus for 1 h before adding to HeLa-hACE2-11 cells. Infection 48 h later was measured by luciferase reporter gene expression. Data are mean ± SD, N = 3 independent replicates. **(D-E)** Authentic BA.1 omicron virus (isolate USA/MD-HP20874/2021) was incubated with sACE2_2_-IgG1 **(D)** or sACE2_2_.v2.4-IgG1 **(E)** for 1 h and added to Calu-3 cells. Infection 48 h later was measured by RT-qPCR for the viral N gene. 3 μM remdesivir (black columns) is a positive neutralization control. Data are mean ± SD, N = 4 independent replicates.

The binding of dimeric sACE2_2_.v2.4-IgG1 to cells expressing S of BA.1 omicron was compared to four monoclonal antibodies authorized for clinical use (**Figure 3B**). Whereas the decoy receptor bound to BA.1 omicron S at low nanomolar concentrations, no substantial binding was observed for REGN10933, REGN10987, or LY-CoV555. Of the tested antibodies, only VIR-7831 bound (consistent with prior reports (Cao *et al*, 2022; VanBlargan *et al*, 2022; Planas *et al*, 2022; Ikemura *et al*, 2022)), albeit less tightly than the engineered decoy in this assay.

HeLa cells expressing human ACE2 were infected with a BA.1 omicron pseudovirus that contains a luciferase reporter gene. Engineered sACE2_2_.v2.4-IgG1 (IC_50_ 18 ± 7 pM, based on the concentration of monomeric subunits) was over an order of magnitude more potent than wild type sACE2_2_-IgG1 (IC_50_ 580 ± 70 pM) (**Figure 3C**), consistent with previous neutralization studies of other SARS-CoV-2 variants. We further tested neutralization of authentic BA.1 omicron virus infecting Calu-3 cells. Based on quantitative measurements of viral RNA, we estimated the IC_50_ for wild type sACE2_2_-IgG1 (**Figure 3D**) and engineered sACE2_2_.v2.4-IgG1 (**Figure 3E**) to be 7.5 ± 9.2 nM and 0.14 ± 0.22 nM, respectively. We conclude that sACE2_2_.v2.4-IgG1 remains exceptionally effective at neutralizing BA.1 omicron.

### The engineered decoy binds tightly to S of BA.2 omicron

The S sequences of BA.1 and BA.2 are separated by 26 mutations and may therefore differ substantially in their interactions with binding proteins. Using flow cytometry to measure binding of monomeric sACE2(18-615) proteins to BA.2 S-expressing cells, it was observed that engineered decoys carrying the v2.4 or CDY14 mutations bound substantially tighter than wild type sACE2 (**Figure 4A**). Furthermore, avid binding of sACE2_2_.v2.4-IgG1 to BA.2 omicron S-expressing cells outperformed the four monoclonal antibodies authorized for emergency use as therapeutics (**Figure 4B**). Importantly, we observed substantially diminished binding of VIR-7831 to BA.2 omicron, in agreement with recent preprints (Zhou *et al*, 2022; Iketani *et al*, 2022).

**Figure 4.**
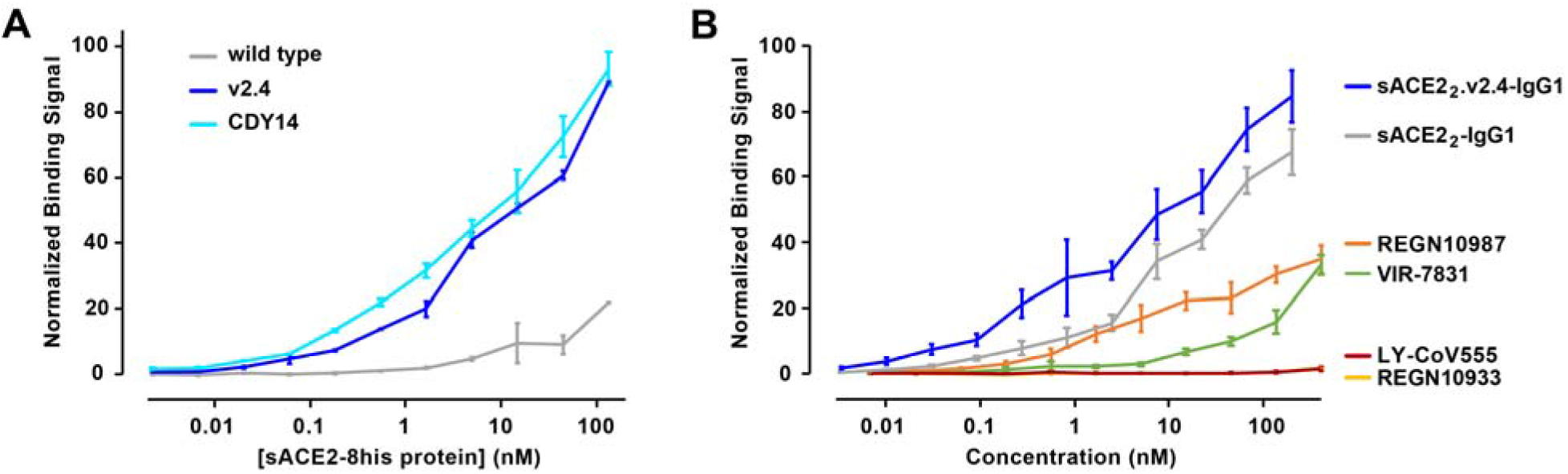
The engineered decoy tightly binds S of BA.2 omicron. **(A)** Using flow cytometry, binding to Expi293F cells expressing S of BA.2 omicron was measured for monomeric sACE2-8his proteins. Data are mean ± SEM, N = 3 independent experiments. **(B)** Binding of antibodies and dimeric sACE2_2_-IgG1 proteins to Expi293F cells expressing BA.2 omicron S measured by flow cytometry. Data are mean ± SEM, N = 4 independent experiments.

### Molecular mechanism of affinity enhancement by the engineered decoy

To understand why ACE2-based decoys carrying the v2.4 mutations bind much tighter to omicron S than wild type ACE2, we modeled the interacting proteins. The cryo-electron microscopy structure of BA.1 omicron RBD bound to ACE2 (PDB 7WPB, 2.79 Å resolution) was used as a template for modeling ACE2.v2.4 bound to both BA.1 and BA.2 RBDs (Yin *et al*, 2022). Structures were relaxed using the ROSETTA energy function (Leman *et al*, 2020). BA.1 and BA.2 omicron RBDs have identical residues at the interface except at position 496 (serine in BA.1 and glycine in BA.2), which is 7.6 Å from ACE2-D38 (Cα-Cα distance). Due to their close similarity at the interface, we describe here only the BA.1 omicron models. All models are provided in online Supporting Information.

Substitution T27Y in ACE2.v2.4 brings the aromatic ring of tyrosine-27 into a cluster of hydrophobic residues on omicron formed by RBD-F456, Y473, A475, and Y489 (**Figure 5**). This is associated with minor backbone movements of RBD loop 1 (a.a. 455-491) and a shift of RBD-Y473 to resolve a small steric clash with the larger ACE2-T27Y side chain. The two other v2.4 mutations, ACE2-L79T and ACE2-N330Y, are at the interface periphery (**Figure 5**). RBD-F486 makes contacts to ACE2-L79T, while RBD-T497 of RBD loop 2 (a.a. 496-506) moves closer to pack against ACE2-N330Y. New polar contacts between the side chain hydroxyls of ACE2-T27Y and RBD-Y473 and between the hydroxyl of ACE2-N330Y and backbone carbonyl of RBD-P499 are also observed, consistent with previous molecular dynamics-based modeling of ACE2.v2.4 bound to the RBDs of SARS-CoV-2 Wuhan, delta, and gamma variants (Zhang *et al*, 2022). We note that the new contacts formed by the ACE2.v2.4 mutations are to RBD residues in loops. Dynamic flexibility of RBD loops to accommodate mutations on the ACE2 surface can help explain why the engineered decoy is broadly active against diverse SARS-CoV-2 variants.

**Figure 5.**
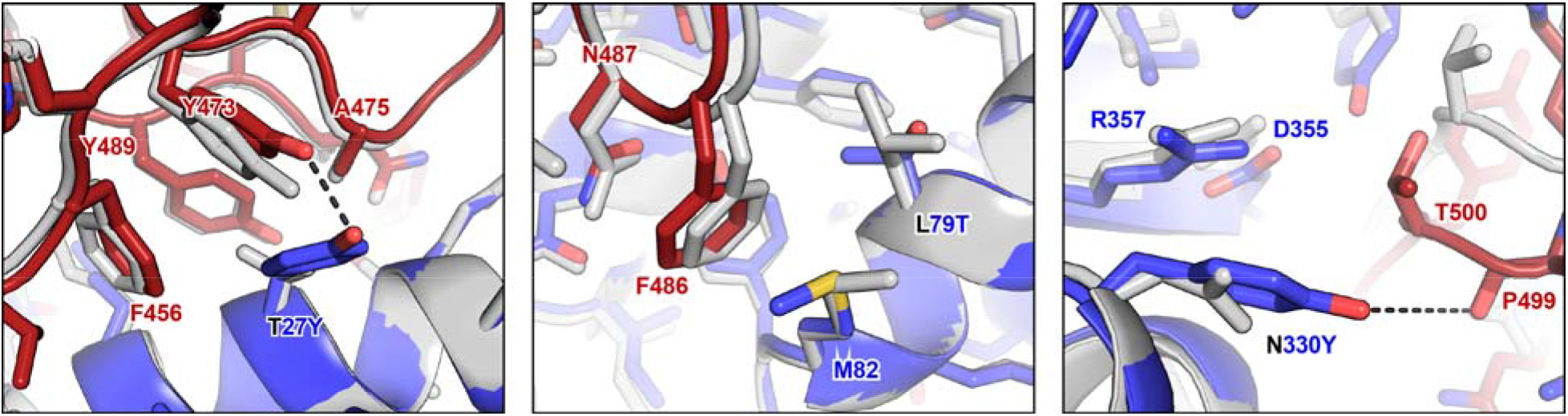
Molecular basis for enhanced affinity of the engineered decoy for omicron RBD. The BA.1 omicron RBD (dark red) bound to wild type ACE2 (grey) and engineered ACE2.v2.4 (blue) was modeled using ROSETTA. Superpositions of the models are shown in the regions surrounding ACE2.v2.4 mutations T27Y (*right*), L79T (*middle*), and N330Y (*right*). New polar contacts formed by the ACE2.v2.4 mutations are indicated with dashed black lines.

## DISCUSSION

The future of the SARS-CoV-2 pandemic is uncertain, but based on the history of the past 2 years, it is expected that new virus variants will continue to emerge as SARS-CoV-2 becomes endemic. There will likely be a continuing need for effective therapeutics, especially as immunity wanes, vaccine hesitancy remains high, and new virus variants emerge that partially escape natural and vaccine-induced antibodies.

Monoclonal antibodies have been important drugs in the clinic and can be co-administered with small molecule drugs that target other features of the SARS-CoV-2 replication cycle. Alarmingly, omicron variants have accumulated enough mutations to partially or fully escape many anti-S antibodies, including VIR-7831 based on our binding data. It is unclear if constant monoclonal antibody development is a viable long-term strategy as new variants continue to emerge. We show here that decoy receptors remain highly potent against both omicron variants and based on their similarity to the native ACE2 receptor, decoys will likely remain effective against future variants as SARS-CoV-2 evolves.

We also addressed the important question of whether sACE2 has additional therapeutic benefits in a SARS-CoV-2 infection beyond the direct binding of the viral spike protein. There has been disagreement in the literature as to whether sACE2-catalyzed turnover of vasoconstrictive and pro-inflammatory peptides will confer therapeutic benefit or whether it is a safety liability. Many groups knocked out catalytic activity when developing candidate decoy receptors, negating ill-defined risks of adverse hypotension (Tanaka *et al*, 2021; Lei *et al*, 2020; Iwanaga *et al*, 2020; Cohen-Dvashi *et al*, 2020; Glasgow *et al*, 2020; Sims *et al*, 2021; Chen *et al*, 2021; Higuchi *et al*, 2021). However, we show here that catalytically inactivated sACE2_2_.v2.4(NN)-IgG1 is not as effective at prolonging survival of hACE2 transgenic mice infected with a lethal virus dose, suggesting that the catalytic activity of ACE2 present in the decoy confers additional therapeutic benefits. We speculate that while neutralizing antibodies are most effective when administered early to patients with mild-to-moderate disease, decoy receptors may have broader reach into hospitalized patient groups due to both neutralizing and ACE2 catalytic activities.

We previously tested IV administered sACE2_2_.v2.4-IgG1 in prophylactic and therapeutic regimens in K18-hACE2 mice, finding 50-100% of mice survived lethal doses of original and gamma viruses (Zhang *et al*, 2022). We now show the protein effectively delays death when inhaled, a mode of delivery that has significant clinical relevance. Inhalation can be readily administered in an outpatient setting and would help reduce the need for in-hospital treatment, which is especially important when hospital resources become scarce during COVID-19 ‘surges’. Inhalation may be the first-line treatment in outpatients with early infection, whereas IV delivery could be reserved for hospitalized patients in which the virus has spread beyond the lungs.

The studies here strengthen the concept of ACE2-based decoy receptors as broadly effective neutralizing agents for SARS-CoV-2 variants with multiple therapeutic mechanisms. Next generation sACE2 decoys with enhanced S affinity and neutralization potency are promising drug candidates for treating an ever-evolving threat long into the future.

## METHODS

### Cell Lines

Expi293F cells (Thermo Fisher) were cultured in Expi293 Expression Medium (Thermo Fisher), 37 °C, 125 r.p.m., 8% CO_2_. HeLa-hACE2-11 (a stable human ACE2 HeLa clone) were grown in Dulbecco’s Modified Eagle’s Medium (DMEM) high glucose (4500 mg/l) with 10% fetal bovine serum (FBS), 100 units/ml penicillin, and 100 μg/ml streptomycin at 37 °C, 5% CO_2_. Calu-3 (ATCC HTB-55) cells were grown in Modified Eagle’s Medium high glucose (4500 mg/l) with 10% FBS, 4 mM L-Glutamine, 1 mM sodium pyruvate, 100 units/ml penicillin, and 100 μg/ml streptomycin at 37 °C with 5% CO_2_. ExpiCHO-S cells (Thermo Fisher) were cultured in ExpiCHO Expression Medium (Thermo Fisher) at 37◻°C, 125◻r.p.m., 8% CO_2_. Vero E6 (CRL-1586, American Type Culture Collection) were cultured at 37 °C, 5% CO_2_, in DMEM supplemented with 10% FBS, 1 mM sodium pyruvate, 1× non-essential amino acids, 100 units/ml penicillin, and 100 μg/ml streptomycin.

### Expression of Proteins

All genes were cloned into the NheI-XhoI sites of pcDNA3.1(+) (Invitrogen) with a consensus Kozak sequence (GCCACC) upstream of the start ATG. Plasmids for sACE2_2_-IgG1 (Addgene #154104), sACE2_2_.v2.4-IgG1 (#154106), sACE2(18-615)-8his (#149268), sACE2(18-615).v2.4-8his (#149664), and Wuhan RBD-8his (#145145) are available from Addgene. Mutations for the CDY14 decoy receptor were introduced into the wild type sACE2 plasmids using extension overlap PCR and confirmed by Sanger sequencing. Monoclonal antibodies, sACE2-8his proteins, and RBD-8his were expressed in Expi293F cells transfected using Expifectamine (Thermo Fisher) according to the manufacturer’s instructions. Transfection Enhancers 1 (5◻μl per ml of culture) and 2 (50◻μl per ml of culture) were added ~18 h post-transfection and medium was collected on day 6. For larger scale production of sACE2_2_-IgG1 proteins (sufficient for animal studies), plasmids (1,000◻ng per ml of culture) were transfected in ExpiCHO-S cells using ExpiFectamine CHO (Thermo Fisher) according to the manufacturer’s instructions. ExpiFectamine CHO Enhancer (6◻μl per ml of culture) was added ~20◻h post-transfection and the temperature was decreased to 33 °C. At days 1 and 5, ExpiCHO Feed (240◻μl per ml of culture) was added. CO_2_ was decreased over days 9-12 to 5% and medium was collected on days 12-14.

### Purification of sACE2_2_-IgG1 and Monoclonal Antibodies

Expression medium was collected after removal of cells by centrifugation (800 g, 4 °C, 10◻min) and the pH was adjusted to 7.5 by adding 1M Tris base. Medium was centrifuged (15,000 g, 4 °C, 20 min) and incubated for 2 h at 4 °C with 2 ml KanCapA resin (Kaneka Corporation) per 100◻ml. Resin was collected in a chromatography column, washed with 10 column volumes (CV) of Dulbecco’s phosphate-buffered saline (PBS), and protein eluted with 4 CV 60LmM sodium acetate (pH 3.7) into 2 CV 1M Tris (pH 8.0). The pH was raised to 7-8 by 1-2 CV 1M Tris base. Eluates were concentrated by centrifugal filtration and separated by size exclusion chromatography using PBS as the running buffer. Peak fractions were pooled, concentrated, and aliquots were stored at −80 °C after snap freezing in liquid nitrogen. Concentrations were determined by absorbance at 280 nm using calculated molar extinction coefficients. For consistency, all concentrations in this manuscript are based on monomeric sACE2 subunits or a H+L chain for antibodies (i.e. concentrations can be considered a measure of binding sites).

### Purification of sACE2-8his and RBD-8his

Expression medium was centrifuged twice (800 g, 4 °C, 10◻min, followed by 15,000 g, 4 °C, 20◻min) and supernatants were incubated at 4 °C, 90 minutes, with 1 ml HisPur Ni-NTA Resin (Thermo Fisher) per 100◻ml. Resin was collected in a chromatography column, washed with >20 CV PBS, washed with ~10 CV PBS containing 20 mM imidazole, and proteins eluted with ~15 CV PBS containing 250◻mM imidazole (pH 8.0). Eluates were concentrated by centrifugal filtration and proteins were separated on a Superdex 200 Increase 10/300 GL column (GE Healthcare) with running buffer PBS. Peak fractions at the expected molecular weight were pooled, concentrated, and aliquots were stored at −80 °C after snap freezing in liquid nitrogen. Concentrations were based on absorbance at 280 nm using calculated molar extinction coefficients.

### S Binding Assay

Human codon-optimized genes encoding N-terminal myc-tagged S proteins of BA.1 and BA.2 omicron were synthesized (Integrated DNA Technologies) and cloned into the NheI-XhoI sites of pcDNA3.1(+) (Invitrogen). The S sequences used in this manuscript have the following mutations from the Wuhan reference sequence (GenBank Accession No. YP_009724390). BA.1: A67V, H69del, V70del, T95I, G142D, V143del, V144del, Y145del, N211del, L212I, ins214EPE, G339D, S371L, S373P, S375F, K417N, N440K, G446S, S477N, T478K, E484A, Q493R, G496S, Q498R, N501Y, Y505H, T547K, D614G, H655Y, N679K, P681H, N764K, D796Y, N856K, Q954H, N969K, and L981F. BA.2: T19I, L24del, P25del, P26del, A27S, G142D, V213G, G339D, S371F, S373P, S375F, T376A, D405N, R408S, K417N, N440K, S477N, T478K, E484A, Q493R, Q498R, N501Y, Y505H, D614G, H655Y, N679K, P681H, N764K, D796Y, L849P, Q954H, and N969K. Expi293F cells were transfected using Expifectamine (Thermo Fisher) according to the manufacturer’s directions. After 24-28 h, cells were washed with cold PBS containing 0.2% bovine serum albumin (PBS-BSA) and added to 3-fold serial titrations of the binding proteins in 96-well round-bottomed plates. Plates were incubated on ice for 30 minutes with regular agitation. Cells were washed with PBS-BSA. For assays examining the binding of monomeric sACE2(18-615)-8his, cells were resuspended in PBS-BSA containing 1:150 polyclonal chicken anti-HIS-FITC (Immunology Consultants Laboratory) and 1:300 anti-myc-Alexa 647 (clone 9B11, Cell Signaling Technology). For assays examining the avid binding of dimeric sACE2_2_-IgG1 and monoclonal antibodies, cells were resuspended in PBS-BSA containing 1:150 polyclonal chicken anti-MYC-FITC (Immunology Consultants Laboratory) and 1:300 anti-human IgG-APC (clone HP6017, BioLegend). Plates were incubated for 30 minutes on ice with occasional mixing, washed twice with PBS-BSA, and analyzed on a BD Accuri C6 flow cytometer using CFlow version 1.0.264.15. Cells were gated by forward and side scatter to exclude dead cells and debris, followed by gating of the myc-positive population. Binding data are presented as the mean fluorescence units (FITC for bound 8his proteins and APC for bound IgG1 proteins) with subtraction of background fluorescence from cells incubated without sACE2 proteins. Data were normalized across independent experiments based on the total measured fluorescence in each experiment.

### Biolayer Interferometry (BLI)

sACE2_2_.v2.4-IgG1 and sACE2_2_.v2.4(NN)-IgG1 were diluted in assay buffer (10◻mM HEPES pH 7.6, 150◻mM NaCl, 3◻mM EDTA, 0.05% polysorbate 20, and 0.5% nonfat dry milk) to 100 nM and immobilized for 60 s to anti-human IgG Fc capture biosensors (Sartorius). Sensors were transferred to assay buffer for 30 s to set the baseline, then transferred to Wuhan RBD-8his for 60 s (association) and transferred back to buffer for 300 s (dissociation). Data were collected on an Octet RED96a and analyzed using instrument software (Molecular Devices) with a global fit 1:1 binding model.

### Catalytic Activity Assay

ACE2 activity was measured with the Fluorometric ACE2 Activity Assay Kit (BioVision) according to the manufacturer’s directions. Fluorescence was read on a Biotek Cytation 5 instrument.

### Gamma SARS-CoV-2 virus amplification and quantification

SARS-CoV-2 isolate hCoV-19/Japan/TY7-503/2021 (P.1/gamma) was obtained from BEI Resources (# NR-54982), NIAID, NIH and propagated in Vero E6 cells. Culture supernatant was collected upon observation of cytopathic effects. Cell debris was removed by centrifugation and passing through a 0.22 μm filter. Supernatant was aliquoted and stored at −80 °C. Virus titers were quantitated by a plaque forming assay using Vero E6 cells.

### Inoculation of SARS-CoV-2 gamma variant in K18-hACE2 transgenic mice

Biosafety level 3 (BSL-3) protocols for animal experiments with live SARS-CoV-2 were performed by personnel equipped with powered air-purifying respirators in strict compliance with NH guidelines for humane treatment and approved by the University of Illinois Animal Care & Use Committee (ACC protocol 21-055 and IBC protocol 20-036). Hemizygous K18-hACE2 mice (strain 034860: B6.Cg-Tg(K18-ACE2)2Prlmn/J) were purchased from The Jackson Laboratory. Animals were housed in groups and fed standard chow. Mice (10-16) weeks old were anesthetized by ketamine/xylazine (50/5 mg/kg, IP). Mice were then inoculated intranasally with 1×10^4^ PFU (plaque-forming units) of SARS-CoV-2 gamma variant suspended in 20 μL of sterile PBS.

### Administration of sACE2_2_.v2.4-IgG1 by inhalation

Mice were placed in a pie cage for aerosol delivery (Braintree Scientific, # MPC-3-AERO). The mice were individually separated and the pie cage can hold as many as 11 mice. A MPC Aerosol Medication Nebulizer (Braintree Scientific, # NEB-MED H) aerosolized 7.5 ml sACE2_2_.v2.4-IgG1 (8.3 mg/ml in PBS) in the nebulizer cup and delivered the aerosol to the mice in the pie cage. Inhalation delivery took approximately 25 minutes. sACE2_2_.v2.4-IgG1 was administered 3 times with doses 36 hours apart and starting 12 hours post virus inoculation. PBS was delivered as control. For the number of animals needed to achieve statistically significant results, we conducted an a priori power analysis. We calculated power and sample sizes according to data from pilot experiments, variations within each group of data, and variance similarities between the groups were statistically compared. Animals with sex- and age-matched littermates were randomly included in experiments. No animals were excluded attributed to illness after experiments. Animal experiments were carried out in a blinded fashion whenever feasible.

### mRNA expression measured by quantitative RT-qPCR

Tissues were homogenized in 1 ml Trizol solution (Thermo Fisher, # 15596026). Tissue homogenates were clarified by centrifugation at 10,000 rpm for 5 min and stored at −80 °C. RNA was extracted according to the Trizol protocol. RNA was quantified by Nanodrop 1000 (Thermo Fisher) and reverse transcribed with Superscript III (Invitrogen # 18080093) using random primers. FastStart Universal SYBR Green Master Mix (Thermo Fisher # 4913850001) was used for relative quantification of cDNA on the ViiA 7 Real-Time PCR System (Thermo Fisher). Primer information is included in **Table 1**.

**Table 1.**
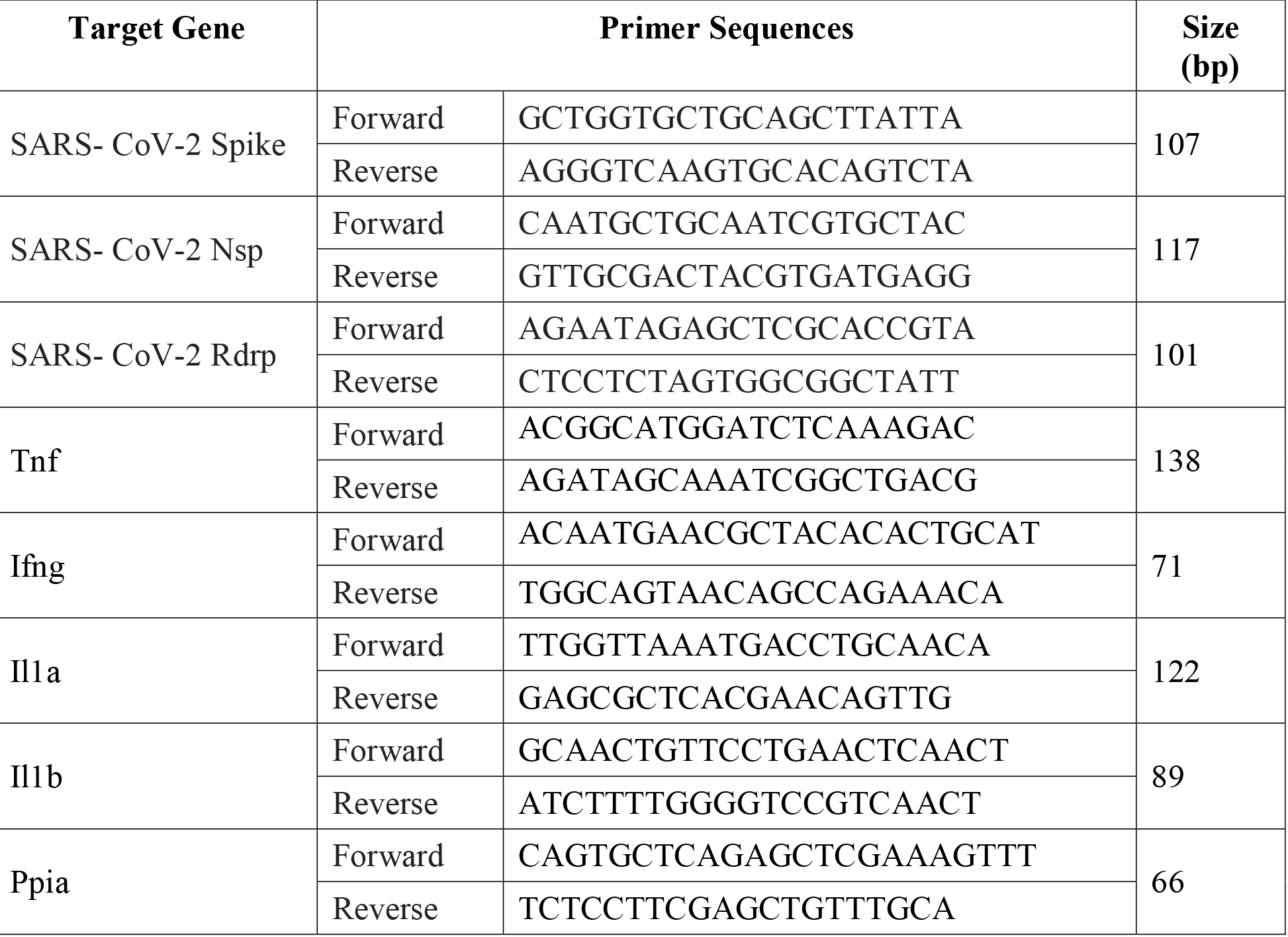
qPCR Primer Sequences

### Histology and imaging

Animals were euthanized before harvesting and fixation of tissues. Lung lobes were fixed with 4% PFA (paraformaldehyde) for 48 hours before further processing. Tissues were embedded in paraffin and sections were stained with hematoxylin and eosin. The slides were scanned by Aperio Brightfield 20x. Images were taken by Aperio ImageScope 12.4.3 and analyzed by Zen software (Zeiss).

### Pseudovirion Production

Pseudoviruses were created using plasmids for SARS-CoV-2 Omicron B.1.1.529 (BA.1) S and HIV-1 proviral vector pNL4-3.Luc.R^−^E^−^ (from the NIH AIDS Research and Reference Reagent Program) containing a luciferase reporter gene. Pseudovirions were created following a polyethylenimine (PEI)-based transient co-transfection on 293T cells. After 5 h, cells were washed with PBS and the medium was replaced with phenol red-free DMEM. 16 h post-transfection, supernatants were collected and filtered through 0.45 μm pore size filter.

### Pseudovirus Inhibition Assay

HeLa-hACE2-11 cells were seeded (5×10^3^ cells/well) onto white-bottomed 96-well tissue culture plates (100 μL/well) and incubated for 16 h, 37 °C, 5% CO_2_. The decoy sACE2 was titrated in SARS-CoV-2 Omicron (B.1.1.529) pseudovirus supernatant and incubated at room temperature for 1 h. The pseudovirus/sACE2 mixtures were added to the target cells. Plates were incubated for 48 h and the degree of viral entry was determined by luminescence using the neolite reporter gene assay system (PerkinElmer). IC_50_ values were determined by fitting dose-response curves with four-parameter logistic regression in GraphPad Prism 8 software.

### Live Omicron Virus Isolation and Neutralization Assay

48 h prior to treatment, 3×10^5^ Calu-3 cells/well were seeded into 24-well plates. Infection was with a clinical isolate of the SARS-CoV-2 omicron BA.1 variant (isolate USA/MD-HP20874/2021) from BEI Resources (NR-56461). Non-treatment controls, 5-fold serial dilutions of decoy sACE2 (final concentrations 500 nM – 0.006 nM), and a high concentration of positive control remdesivir (3 μM) were added to the same volume of SARS-CoV-2 (final MOI = 0.01) and incubated at room temperature for 1 h. Then the mixture was added to the monolayer of cells and incubated 1 h at 37 °C, 5% CO_2_. The mixture was removed, cells washed with PBS, and monolayers overlayed with infection media (2% FBS). After 48 h, 100 μL of cell supernatants were collected and added to 300 μL of TRIzol. RNA was isolated using Invitrogen’s PureLink RNA Mini Kits according to the manufacturer’s protocol. Quantitative RT-qPCR was carried out using 5 μL of RNA template in TaqMan Fast Virus 1-Step Master Mix using primers and probes for the N gene (N1 primers) designed by the U.S. Centers for Disease Control and Prevention (IDT cat# 10006713). A standard curve was generated using dilutions of synthetic RNA from the SARS-CoV-2 amplicon region (BEI Resources, NR-52358). All experiments prior to RNA isolation were performed in a Biosafety Level 3 facility.

### Structural Modeling

Models of SARS-CoV-2 omicron variants BA.1 and BA.2 RBDs bound to ACE2 and ACE2.v2.4 were generated using template PDB 7WPB. 7WPB captures the structure of BA.1 RBD complexed with human ACE2. Models were generated using one round of Rosetta Relax Design protocol with Rosetta score function ref2015 and an appropriate mutation set defined for each complex. Relax Design allows for backbone perturbations for the entire structure, making it ideal for allowing movement beyond mutation sites to better accommodate desired mutants (Conway *et al*, 2014). Following Relax Design, each complex underwent an additional 5 rounds of Rosetta Relax with score function ref2015. Lowest energy models were selected from the resulting 5 decoys per complex.

### Statistics and Reproducibility

Quantification of replicate experiments is presented as mean ± SD or SEM as described in figure legends. Statistical tests are described in figure legends. Based on our experience, we expect changes in the gene/protein expression and function measurements to be detected with 4 mice per group, so the effect size was determined as N = 4 independent mice. The variance between groups that are being statistically compared was similar.

### Study Approval

All aspects of this study were approved by the office of Environmental Health and Safety at University of Illinois at Chicago prior to the initiation of this study. All work with live SARS-CoV-2 was performed in a BSL-3 laboratory by personnel equipped with powered air purifying respirators.

## Supporting information

Model of BA.1 RBD bound to wild type ACE2

Model of BA.1 RBD bound to ACE2.v2.4

Model of BA.2 RBD bound to wild type ACE2

Model of BA.2 RBD bound to ACE2.v2.4

## ACKNOWLEDGEMENTS

This work was supported in part by NIH grants R43-AI162329 to E.P. and K.K.C; R01HL157489 to L.Z.; R01-HL162308 to A.B.M and J.R. The following reagents were obtained through BEI Resources: Isolate hCoV-19/Japan/TY7-503/2021 (P.1; NR-54982) and isolate hCoV-19/USA/MD-HP20874/2021 (BA.1; NR-56461).

## AUTHOR CONTRIBUTIONS

L.Z. performed the mouse study. K.K.N. cloned omicron plasmids, expressed proteins, and tested binding. L.M.C. and L.R. tested pseudovirus and authentic virus neutralization. K.K.C. measured BLI kinetics and purified proteins. C.A.D. examined ACE2 catalytic activity. A.A. modeled structures. K.S. cloned plasmids and purified proteins. E.P. purified proteins. E.P. and L.Z. drafted the manuscript. L.Z., L.R., J.R., A.B.M., and E.P. supervised research and planned experiments. All authors contributed to manuscript edits.

